# Experimental characterization and automatic identification of stridulatory sounds inside wood

**DOI:** 10.1101/2021.09.08.459381

**Authors:** Carol L. Bedoya, Ximena J. Nelson, Eckehard G. Brockerhoff, Stephen Pawson, Michael Hayes

**Affiliations:** School of Biological Sciences, University of Canterbury, Private Bag 4800, Christchurch, New Zealand; Swiss Federal Research Institute WSL, Zürcherstrasse 111 8903, Birmensdorf, Switzerland; SCION (New Zealand Forest Research Institute), PO Box 29237, Christchurch, New Zealand; Department of Electrical and Computer Engineering, University of Canterbury, Private Bag 4800, Christchurch, New Zealand

**Keywords:** Acoustic communication, Bark beetles, Forest insects, Scolytinae, Sound production

## Abstract

The propagation of animal vocalizations in water and in air is a well-studied phenomenon, but sound produced by bark and wood boring insects, which feed and reproduce inside trees, is poorly understood. Often being confined to the dark and chemically-saturated habitat of wood, many bark- and woodborers have developed stridulatory mechanisms to communicate acoustically. Despite their ecological and economic importance and the unusual medium used for acoustic communication, very little is known about sound production in these insects, or their acoustic interactions inside trees. Here, we use bark beetles (Scolytinae) as a model system to study the effects of wooden tissue on the propagation of insect stridulations and propose algorithms for their automatic identification. We characterize distance-dependence of the spectral parameters of stridulatory sounds, propose data-based models for the power decay of the stridulations in both outer and inner bark, provide optimal spectral ranges for stridulation detectability, and develop automatic methods for their detection and identification. We also discuss the acoustic discernibility of species cohabitating the same log. The species tested can be acoustically identified with 99% of accuracy at distances up to 20 cm and detected to the greatest extent in the 2-6 kHz frequency band. Phloem was a better medium for sound transmission than bark.

## 1. BACKGROUND

Forest soundscapes are a recurrent topic in acoustic, ecological, and sociological studies (Dumyahn and Pijanowski, 2011; Ross and Mason, 2017; Burivalova *et al*., 2019). These sounds can inform our understanding of the interactions between animals and their habitat (Dumyahn and Pijanowski, 2011; Pijanowski *et al*., 2011). Nonetheless, attention is biased toward sounds that propagate through air or water - neglecting local soundscapes occurring in other propagation media. One of these is wood, within which some insects (e.g., bark beetles (Scolytinae), wood borers (e.g., Cerambycidae, Bostrichidae and Ptinidae), pinhole borers (Platypodinae), and termites (Isoptera)) communicate acoustically (Birch and Keenlyside, 1991; Lai *et al*., 2017; Bedoya *et al*., 2021). We know very little about communicatory interactions inside wood/bark and the transmission of acoustic information within these media (Hill *et al*., 2019). In order to address this, we use bark beetles (Coleoptera: Curculionidae: Scolytinae) to study the propagation and attenuation of stridulatory sounds inside trees. We also propose strategies for the automatic acoustic detection and identification of bark beetles and woodborers so that they can be acoustically studied without disrupting their natural habitat.

Bark beetles are a subfamily of weevils that spend most of their life cycle inside plant tissue (Kirkendall *et al*., 2015; Raffa *et al*., 2015) and are one of the very few animals that have evolved sound production mechanisms to communicate inside plants (Bedoya *et al*. 2021; Hofstetter *et al*., 2019). Although ‘bark beetle’ is usually used to refer to all the Scolytinae, ‘true bark beetles’ are the subset that live, feed, and reproduce in the phloem tissue of trees (i.e., phloeophagy) (Kirkendall, 1983; Wood and Bright, 1982). There are ca. 6000 described species of Scolytinae (Kirkendall *et al*., 2015) distributed in all regions of the world except Antarctica (Raffa *et al*., 2015). Previous studies of bark beetle life history and behavior typically focus on the <1% of species that are important forest pests that attack and potentially kill trees (Grégoire *et al*., 2015; Kirkendall *et al*., 2015).

Bark beetles typically construct an intricate system of tunnels (also referred to as galleries) within trees, where adults and larvae feed and complete their development (Hofstetter *et al*., 2019). Some bark beetles use airborne pheromones to communicate over large distances that facilitate aggregation or disrupt aggregations of conspecifics (Raffa *et al*., 2015), and acoustic signals, on and within the host, for intraspecific communication over short distances (Rudinsky and Michael, 1973; Ryker and Rudinsky, 1976). However, the sounds of only a few, typically economically important, species have been reported in the literature. From the limited data available, acoustic signaling appears to be widespread within the group, but remains poorly documented (Barr, 1969; Lyal and King, 1996; Bedoya *et al*., 2021). Sound production in Scolytinae is mediated by three predominant types of stridulatory organs: elytro-tergal, vertex-pronotal, and gula-prosternal (Barr, 1969; Lyal and King, 1996; Bedoya *et al*., 2021). These organs can arise in one, both, or neither of the sexes, and, in studies to date where both sexes stridulate, the organ and the signals are usually sexually dimorphic (Lyal and King, 1996; Bedoya *et al*., 2021; Hofstetter *et al*., 2019). Each stridulatory organ consists of two parts; A) a static file of teeth, also known as pars stridens, and B) a movable plectrum consisting of a set of spines, tubercles, or teeth that rubs against the static file (Barr, 1969). Acoustic characteristics of the stridulatory sounds vary between species (Fleming *et al*., 2013; Yturralde and Hofstetter, 2015; Bedoya *et al*., 2019a). Such characteristics are also dependent on the behavioral context (Fleming *et al*., 2013; Bedoya *et al*., 2019a), as acoustic communication is used in several functions, including distress, pre-mating recognition, rivalry, and copulation (Barr, 1969; Lyal and King, 1996; Fleming *et al*., 2013).

Given how little we know about bark beetle and woodborer stridulatory behavior in in general, it is unsurprising that the effect of the propagation medium on their sounds has not been assessed. Previous studies have mostly focused on the analysis of mechanical sounds (e.g., chewing), or the vibrational movement of insect larvae and pupae (Mankin *et al*., 2011; Jalinas *et al*., 2019; Sutin *et al*., 2019). However, the specific effect of wood and bark tissue on the propagation of acoustic communication (i.e., signals produced by acoustic organs) has yet to be investigated. Several theoretical methods have been developed for studying sound attenuation and absorption by trees (Burns, 1979; Price *et al*., 1988, Li *et al*., 2020) and wood (Wassilieff, 1996; Legg and Bradley, 2016); however, these models are typically used to estimate wood characteristics and have yet to be experimentally verified using biotic sound sources.

The goal of our study was to address several unresolved issues related to the propagation of stridulatory sounds inside wood, so that this new information can be used for the acoustic detection and identification of insects inside trees. We analyzed the acoustic signals of two bark beetles, *Hylastes ater* Paykull and *Hylurgus ligniperda* (Fabricius), in order to characterize distance-dependent changes in the spectro-temporal features of stridulations propagating through wood. We determine which part of the audible spectrum is the most suitable to acoustically detect stridulations and investigate the maximum distances at which the presence of a bark beetle can be acoustically detected and the species identified. Then, we propose a data-based model for the attenuation of stridulatory sounds through wood, taking into consideration the type of tissue and its width. Finally, we implement a method for the acoustic detection and identification of stridulations, and provide suggestions for future improvements.

## 2. METHODS

### 2.1. Subjects

*Hylastes ater* and *Hylurgus ligniperda* were selected to study the propagation of stridulatory sounds inside wood because physical interactions (e.g., touching) trigger stridulatory behavior in males (Bedoya *et al*., 2021), and thus, sound production can be manually elicited by the researcher. *Hylurgus ligniperda* has one of the highest known calling rates of all bark beetles (Bedoya *et al*., 2019a, 2021) and tends to sing uninterruptedly for long periods (i.e., tens of minutes). *Hylastes ater* also responds acoustically to physical stimulation, although the duration of the stridulatory behavior is shorter than in *H. ligniperda*. Both species are less than 6 mm in body length (*H. ater* 4.0 mm and *H. ligniperda* 5.0 mm, on average; Fig. 1) and colonize a variety of conifers, but mainly *Pinus* spp., including economically important species (Brockerhoff *et al*., 2003). Insects were manually collected from recently felled *Pinus radiata* D.Don logs in Bottle Lake Forest, Christchurch, New Zealand (−43°27’8.64” S 172°41’42.00” E).

**FIG. 1.**
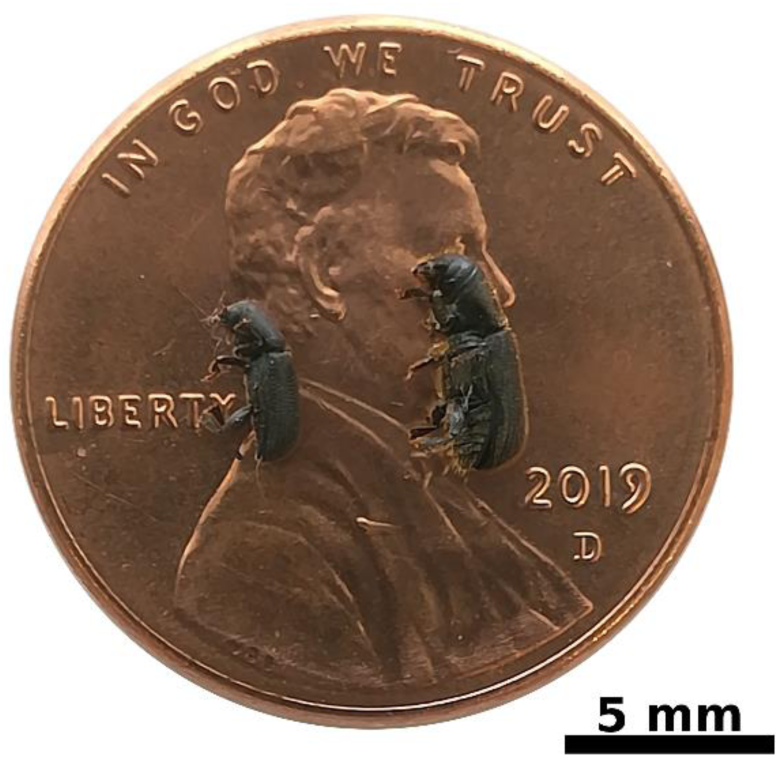
Size comparison of males of *Hylastes ater* (left) and *Hylurgus ligniperda* (right) on an American penny.

### 2.2. Experimental setup

Recordings were acquired using a 352A24 monoaxial accelerometer (PCB piezotronics, Depew, USA) and a 744T recorder (Sound Devices, Reedsburg, USA). Analyzed signals were of one minute duration at a sampling frequency of 44100 kHz, 48 dB gain, and 24-bit resolution. Two *P. radiata* logs (200 cm long, with respective mean±SD diameters of 19.2 ±0.3 and 26.4 ±0.7 cm) were used during the experiment. The logs were held inside a temperature-controlled room at a constant temperature of 23°C for the duration of the experiments (14 days). Humidity inside the phloem was measured using a SHT85 sensor (Sensirion, Stäfa, Switzerland) after collecting data from each individual in each log in order to ensure humidity did not decline substantially.

### 2.3. Data collection

To estimate the effect of the outermost bark layer on signal acquisition, the experimental procedure was performed on the bark surface and inside the phloem tissue of two *P. radiata* logs of different diameters. Most bark- and woodborers live underneath the outermost bark tissue; thus, whether to pierce the bark is an important question that naturally arises before performing acoustic data acquisition in trees. Therefore, we tested tissue effects on each log, which had different average thicknesses of bark (3.3±1.2 mm; 8.5±1.3 mm) and phloem (2.6±0.5 mm; 3.1±0.8 mm). Subsequently, we recorded acoustic signals of *H. ligniperda* and *H. ater* at nine pre-allocated distances (5, 10, 15, 20, 30, 40, 60, 80, and 100 cm) from the position of the stridulating beetle (Fig. 2). Five beetles of each species were individually recorded in both logs with sensors located on the bark and in the phloem tissue at the respective nine distances (N=5 per species, 9 distances, 3 factors (beetle species, tissue type, log thickness), 2 levels per factor, n=45 per treatment, 360 recordings in total).

**FIG. 2.**
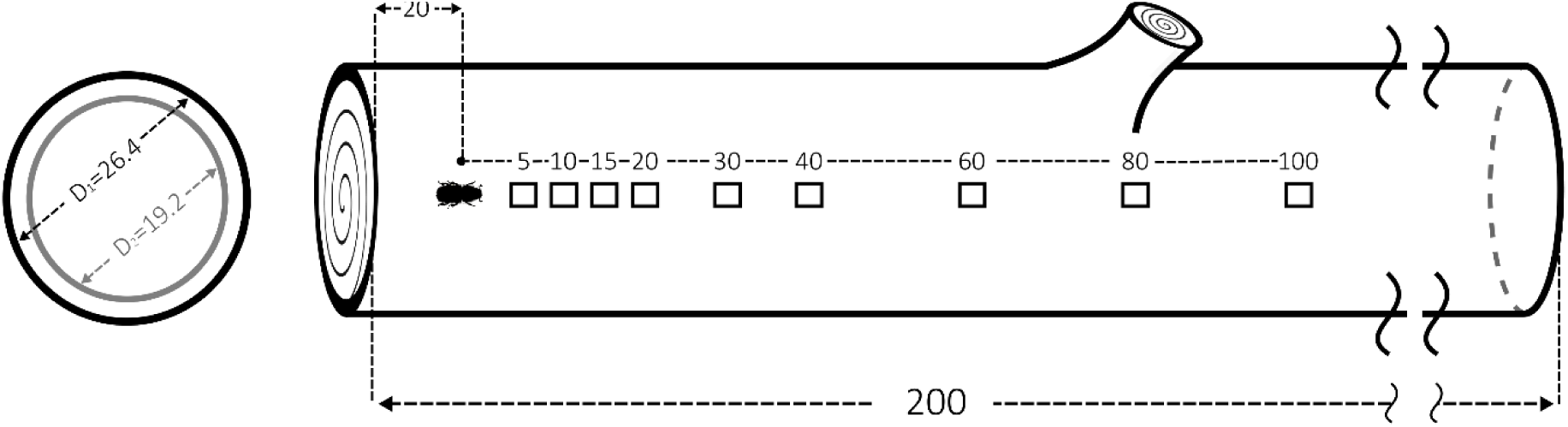
Experimental setup for the analysis of sound propagation of two bark beetle species (*Hylurgus ligniperda* and *Hylastes ater*) in wood. D_1_ and D_2_ are the average diameters of the *Pinus radiata* logs used for testing. The beetle was placed 20 cm from one end of the log. Stridulatory sounds produced by the individual were recorded at nonlinearly-spaced distances from 5 to 100 cm. This procedure was repeated in the bark and phloem layer of each log. Dimensions in cm (not to scale).

Beetles were individually inserted into a pre-drilled hole (0.5 cm diameter) through the outer bark into the phloem, at a distance of 20 cm from the edge of the log (Fig. 2). Then, the elytra of each beetle was softly touched with a paintbrush (Bockingford, 5700R, size 1) to trigger sound production as per Bedoya *et al*., (2019a,c). To record stridulations, the vibrational sensor (accelerometer) was attached to the bark, along the grain, using Blu-Tack™ at any of the nine discrete distances from 5 to 100 cm (Fig. 2). Once the signal was acquired, the sensor was randomly moved to a different position and data collection started again until data were acquired from all nine pre-allocated distances for each beetle. After signals were recorded on the bark of each of the two logs, 1 cm^2^ holes were carved into the bark until the phloem tissue was accessible. Then, the sensor was placed on the phloem and the experimental procedure was repeated, as previously described on the bark, using the same individuals.

### 2.4. Data analysis

#### 2.4.1. Spectrogram and power spectrum estimation

Spectrograms used for visualization were generated using a FFT of 1024 bins and a symmetric flat top window of 1024 samples with 3/4 overlap. Plots of power spectral density (PSD) were generated by averaging the PSDs of all test subjects for each species at each of the analyzed distances. The frequency-dependent power decay was estimated by averaging the mean PSD values from the signals of each species at every pre-determined distance (5-100 cm) in frequency bands of 2 kHz. Power spectral densities were estimated using Welch’s method. Power values hereafter are shown in dB Full Scale (dBFS), using the maximum power value of all signals as reference for the scaling.

#### 2.4.2. Experimental models

Attenuation models were generated by fitting the average power decay from individuals of *H. ligniperda* and *H. ater* to exponential functions 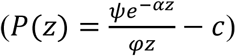, where *α* is the frequency-dependent attenuation coefficient, *z* is the distance between the sensor and the source, and *φ, ψ*, and *c* are model constants in dB. The exponential fitting was performed using nonlinear least squares on the averaged power levels at each distance. Recordings were separated by species, type of tissue, and tissue width, and were modeled independently. The relationship between the tissue width and the attenuation coefficient was determined *a posteriori* by fitting a linear model between the values of both parameters. The root mean square error was estimated as measure of goodness of fit, and 95% confidence intervals were shown for the estimated parameters.

#### 2.4.3. Automatic acoustic detection and identification

Since acoustic features are dependent on distance from the source, we implemented several supervised and unsupervised automatic acoustic detection methods to determine the maximum distance at which species can be reliably identified. We extracted all the stridulations from our recordings using an energy-based segmenter and estimated five acoustic features for each of them (centroid frequency, dominant frequency, bandwidth, duration, and mean amplitude). Then, we used four different clustering algorithms and seven classification techniques to estimate the accuracy of the species identification.

##### 2.4.3.1. Segmentation and feature extraction

Stridulations were segmented from the spectrogram using a threshold-based approach (Bedoya *et al*., 2019a). The method consisted of averaging the values of the spectrogram in the time domain, and using the mean value of this new vector as a threshold for identifying the start and end of a stridulation. Five acoustic features were then estimated for each stridulation: the centroid frequency, dominant frequency, bandwidth, duration, and amplitude. The centroid frequency *f*_*c*_ was estimated using: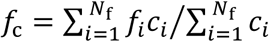, where *c*_*i*_ is the i^th^ value of the mean spectrum, and *f*_*i*_ is the current frequency bin. This frequency is analogous to the centre of mass in mechanical systems (Le et al., 2011). The dominant frequency was the frequency bin with the maximum power value. Bandwidths were determined by the upper and lower cut-off frequencies of the mean spectrum of each call (cut-off 3 dB). Duration was defined as the length of the call. The mean power of the spectrum was used as the amplitude feature. All the acoustic features were normalized (0-1) before using them as input for the clustering and classification algorithms. Since acoustically detecting the presence of bark beetles is possible even if the specific species cannot be discerned, we also estimated average centroid frequencies throughout the log in order to find the distance at which species are spectrally distinguishable.

##### 2.4.3.2. Clustering and Classification

To evaluate the discernibility of species with distance, all stridulations were clustered into two groups using four unsupervised learning techniques (K-means, Fuzzy c-means (FCM), DBSCAN, and Gaussian mixture models (GMM)) applied on the five extracted acoustic features. For the K-means, the squared Euclidean distance was used as metric for minimization. For the FCM, the fuzzy partition matrix exponent that controls the degree of fuzzy overlap (i.e., the fuzzifier) was set to 2. In the GMM case, model likelihood was optimized using the expectation-maximization algorithm. Finally, for DBSCAN, 50 was selected as the minimum number of points and ε=0.25.

All the classification algorithms (i.e., supervised learning) were trained to identify both species using 5-fold cross-validation (80% training - 20% test) at each specific distance. The decision tree used the Gini’s diversity index as split criterion with four as maximum number of splits. Linear and quadratic discriminant analyses used full covariance matrices. The Naive Bayes Classifier was implemented with a Gaussian kernel, while Support Vector Machines (SVMs) were tested with linear, quadratic, cubic, and gaussian kernels. Results for the K-nearest neighbors algorithm (KNN) are presented for Euclidean, Cosine, and Minkowski distances using equal distance weights and 10 neighbors. Decision trees, linear discriminant analyses (LD), and KNNs were also used in ensemble. Bag ensemble was used for the decision tree (number of learners = 30, maximum number of splits = 712), whereas both LD and KNN used subspace ensemble (30 learners, and 3 subspace dimensions).

Accuracies, defined as (Tp+Tn)/(Tp+Tn+Fp+Fn), were reported as general performance measurements for all classification and clustering algorithms. Here, Tp, Tn, Fp, and Fn are the rates of true positives, true negatives, false positives, and false negatives, respectively. Bark beetle acoustic terminology is based on Bedoya *et al*. (2019a). All figures and mathematical models were coded in Matlab 2018b.

## 3. RESULTS

*Hylurgus ligniperda* and *H. ater* possess single-note quasiperiodically-repeating calls that are strongly attenuated by the phloem (Fig. 3). With increasing distance, signal intensity decreases and spectral content (e.g., bandwidth) compresses, while some temporal features (e.g., duration) shrink, and others (e.g., inter-syllable interval) expand due to frequency-dependent attenuation (Wanniarachchi *et al*., 2017) (Fig. 3).

**FIG. 3.**
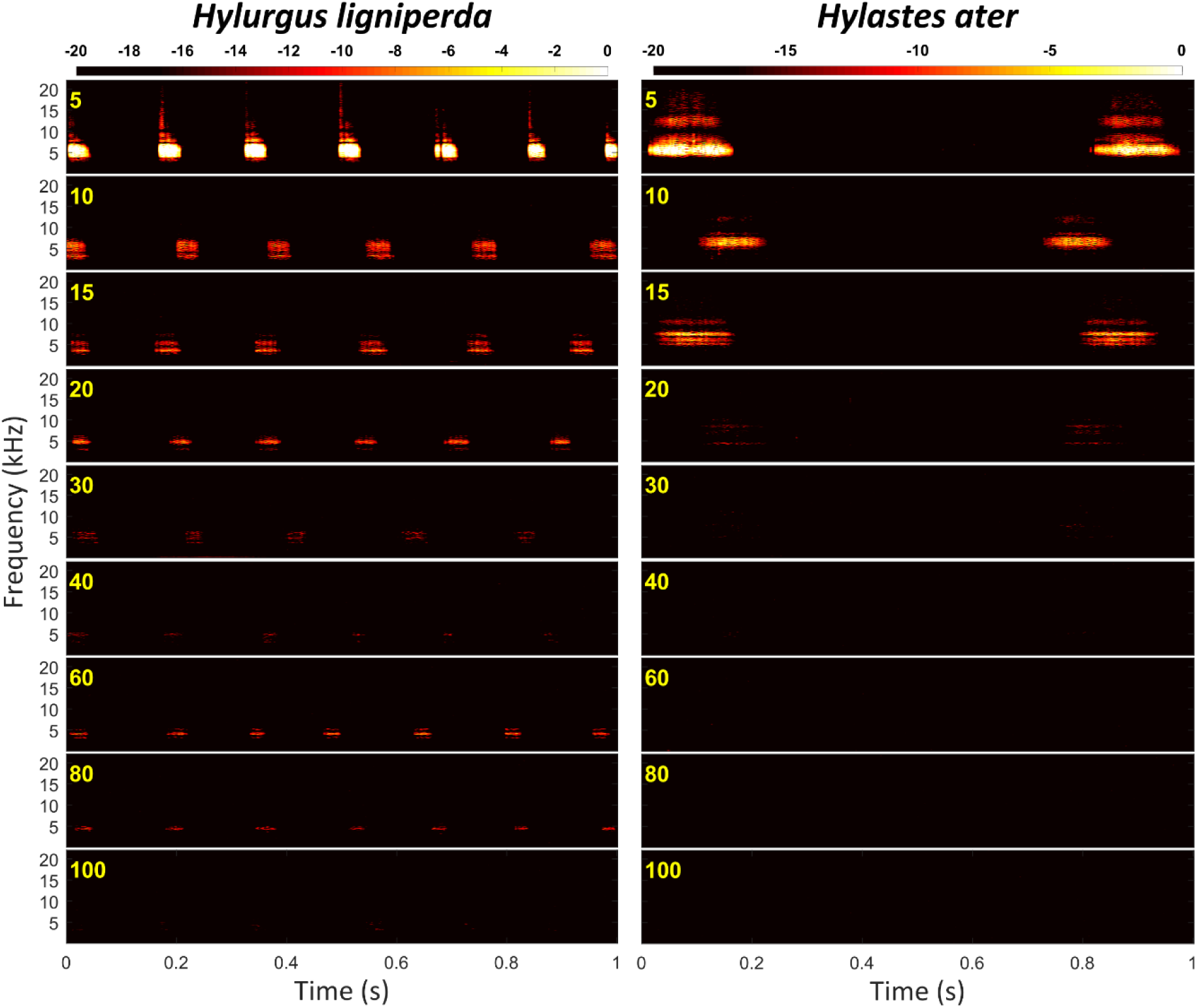
Stridulations of two individuals of *Hylurgus ligniperda* (left) and *Hylastes ater* (right) on *Pinus radiata* phloem recorded from 5 (top) to 100 (bottom) cm from source. Colorbars in dBFS. In some individuals, sounds of *H. ligniperda* are detectable at 100 cm, while those of the smaller *H. ater* are only detectable up to 40 cm.

### 3.1. Power decay

We estimated the power spectra of recordings with *H. ligniperda* and *H. ater* stridulations (Fig. 4). In both species, power was mostly concentrated between 3 and 7 kHz, and decayed with distance. *Hylurgus ligniperda*, the bigger species, had very noticeable power distributions up to 40 cm, whereas *H. ater* had a pronounced decrease in power after 20 cm (Fig. 4).

**FIG. 4.**
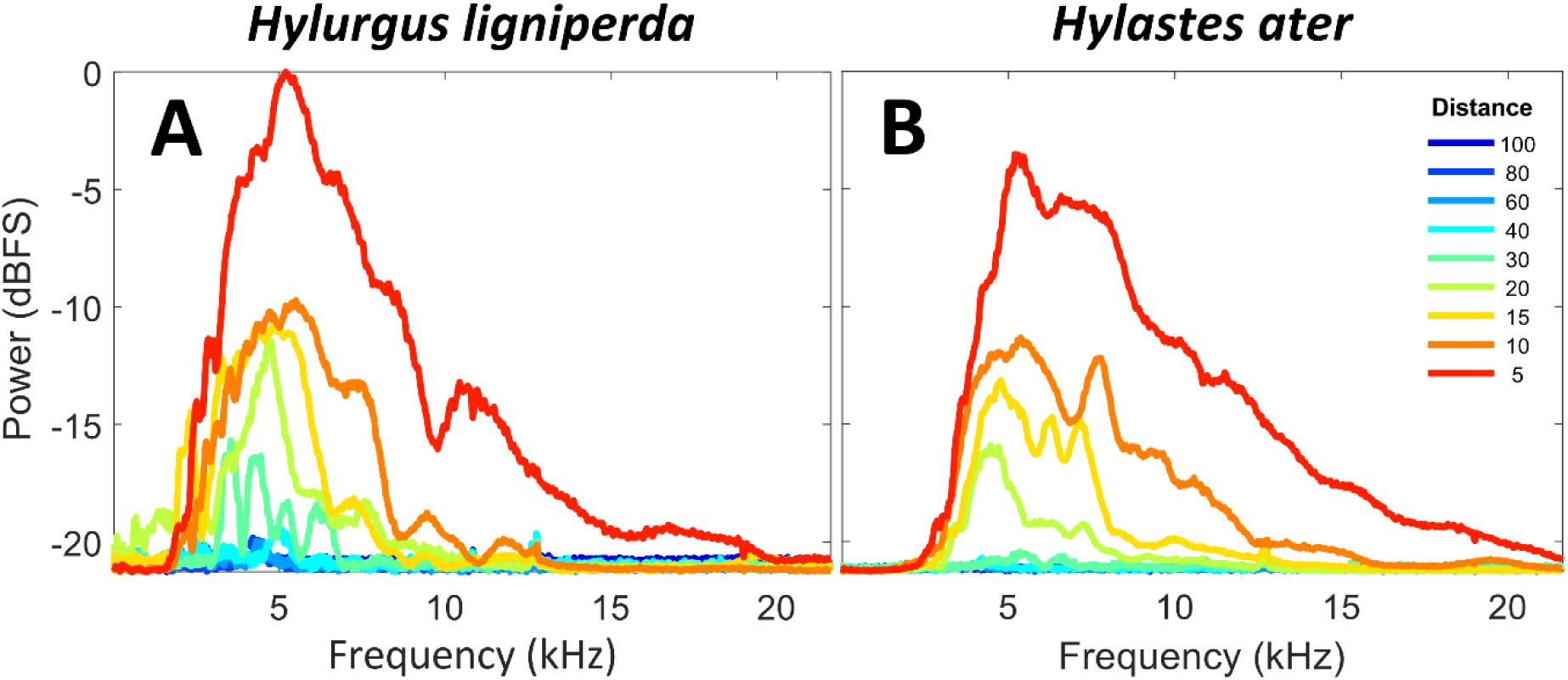
Average power spectral density (PSD) of the stridulatory sounds of (A) *Hylurgus ligniperda* and (B) *Hylastes ater* recorded at distances of 5 to 100 cm from source. Most power is concentrated between 3 and 7 kHz. Averaging was for all individuals of each species, after PSD estimation for the entire dataset (phloem and bark).

In order to localize a specific frequency band for acoustic detection, we divided the spectrum into 2 kHz bands and measured the average power decay at each distance (Fig. 5). The most suitable frequency range for detecting individuals at long distances was 4-6 kHz (Figs. 3, 5). Power decays significantly after 20 cm for *H. ater*, and 40cm for *H. ligniperda* (Fig. 3). After 40cm, sounds are slightly perceptible for some *H. ligniperda* individuals, but only in the 2-6 kHz frequency band (Figs. 3, 5).

**FIG. 5.**
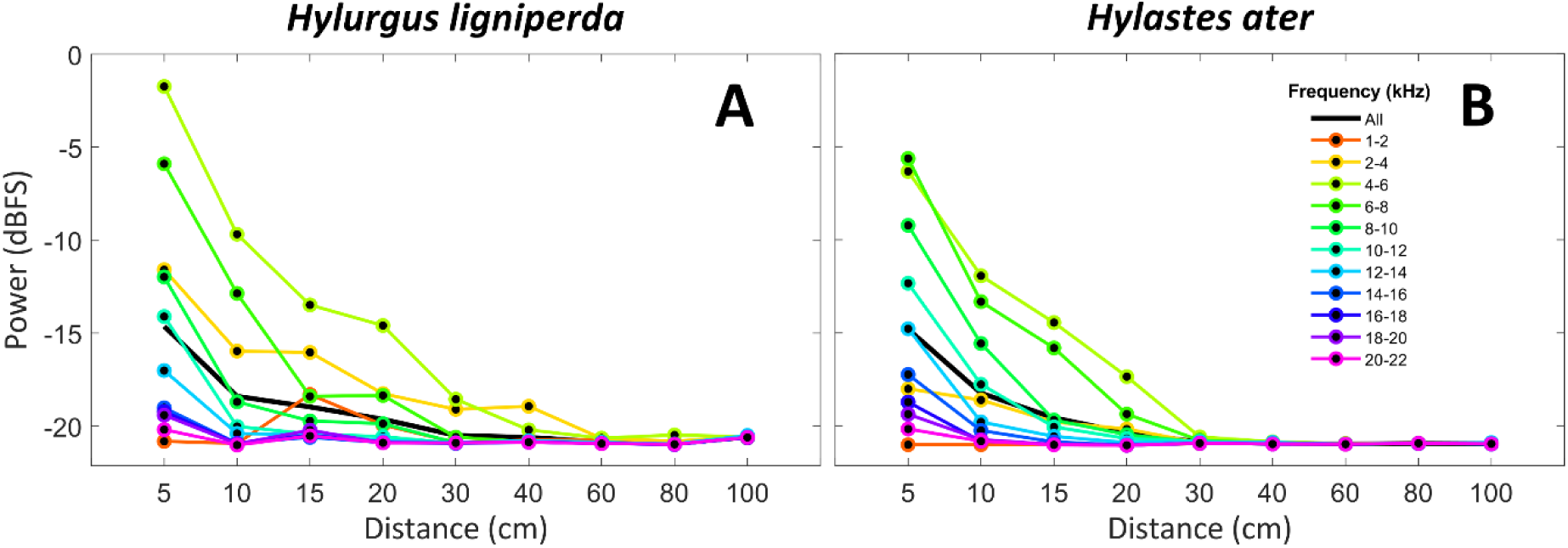
Average power decay with distance of the stridulations of (A) *Hylurgus ligniperda* and (B) *Hylastes ater* estimated in frequency bands of 2 kHz for the entire dataset (phloem and bark). Most stridulations can be detected at their furthest reach using solely the 4-6 kHz frequency band, where spectral components are less attenuated.

### 3.2. Attenuation modelling

Our experimental results show that tissue width significantly reduces power over distance - the wider the tissue (bark/phloem), the more the signal amplitude is attenuated (Fig. 6). Attenuation in bark was stronger than in phloem, and Stridulations of *H. ater* (the smaller species) attenuate faster than those of *H. ligniperda* (Fig. 6, Table I). The phloem is the tissue that transports the soluble organic compounds inside trees; thus, it is more humid and presents less impedance to sound transmission (Yang *et al*., 2015).

**FIG. 6.**
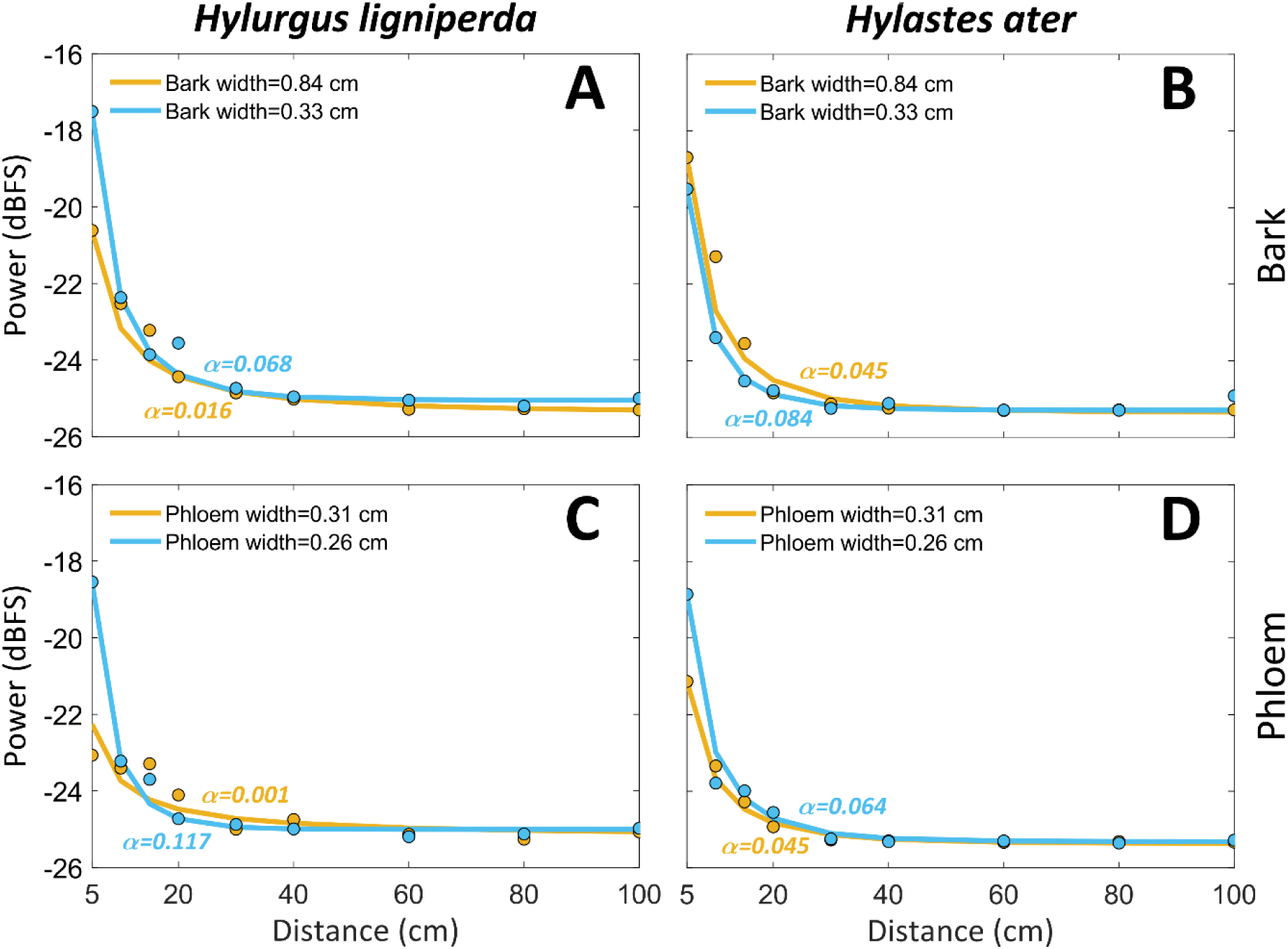
Experimental models for the power attenuation with distance of stridulatory sounds inside *Pinus radiata* logs. α is the attenuation coefficient of an exponentially decaying function (*ψe*^(−α·z)^/*φz* − *c*). Models are shown for two different types of tissue (phloem and bark) of different widths, and two bark beetle species (*Hylurgus ligniperda* and *Hylastes ater*). Data points represent the average power for all the individuals at that distance.

**TABLE 1.** Parameters of the experimental models for the power decay with distance of stridulatory sounds inside *Pinus radiata* bark and phloem. Power levels at each distance were fitted to an exponentially decaying function *P*(*z*) = *ψe*^(−α·z)^/*φz* − *c*, where *z* is the distance in cm and α is the attenuation coefficient. The root mean square error (RMSE) is shown as measure of goodness of fit.

| Species | Tissue | Width (cm) | $\alpha$ | $\psi$ | $\varphi$ | c | RMSE |
| --- | --- | --- | --- | --- | --- | --- | --- |
| *H. ligniperda* | Bark | 0.84 | 0.016 | 0.682 | 0.026 | 25.36 | 0.3021 |
| *H. ligniperda* | Bark | 0.33 | 0.068 | 0.512 | 0.009 | 25.05 | 0.2701 |
| *H. ligniperda* | Phloem | 0.31 | 0.001 | 0.922 | 0.066 | 25.20 | 0.6128 |
| *H. ligniperda* | Phloem | 0.26 | 0.117 | 0.796 | 0.014 | 25.01 | 0.3051 |
| *H. ater* | Bark | 0.84 | 0.045 | 3.359 | 0.080 | 25.34 | 0.6787 |
| *H. ater* | Bark | 0.33 | 0.084 | 1.062 | 0.024 | 25.28 | 0.1846 |
| *H. ater* | Phloem | 0.31 | 0.045 | 0.900 | 0.034 | 25.36 | 0.1917 |
| *H. ater* | Phloem | 0.26 | 0.064 | 1.044 | 0.023 | 25.31 | 0.3857 |

Aside from the exponential models, we generated linear models to correlate our attenuation coefficients (α) with the width of the tissue:

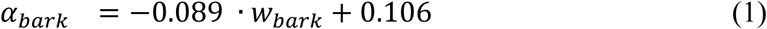

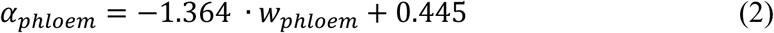

where *α*_*bark*_ and *α*_*phloem*_ are the attenuation coefficients (dB/cm) for bark and phloem, depending on the width of the bark (*w*_*bark*_) and the phloem (*w*_*phloem*_), respectively, with widths in cm.

### 3.3. Species identification

Our data showed that, for *H. ater*, the centroid frequency stabilizes at 30 cm, and that beyond 40 cm, the two species were indistinguishable using solely spectral content (Fig. 7).

**FIG. 7.**
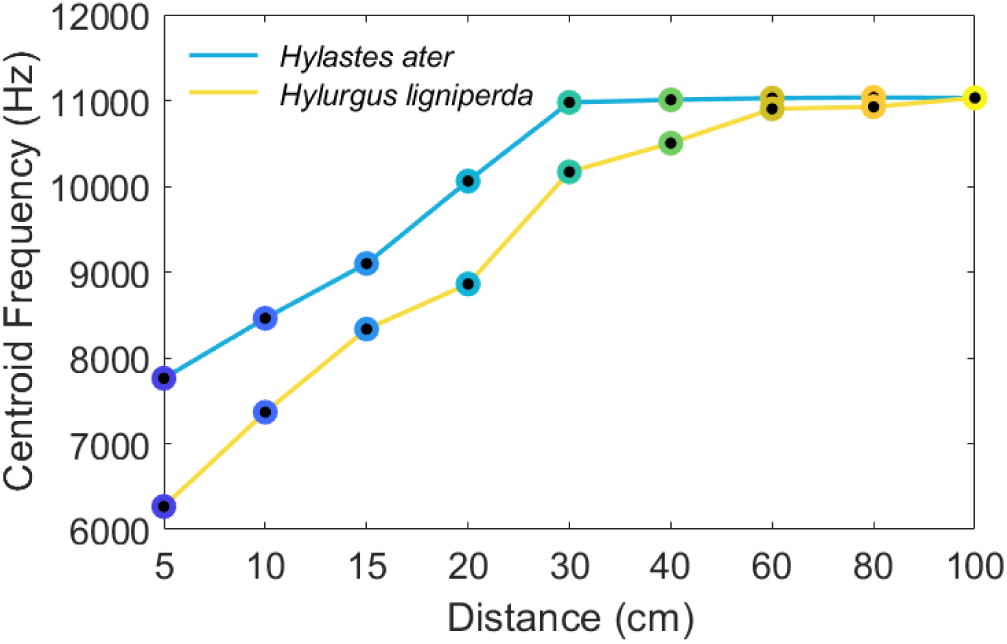
Centroid frequency of *Hylurgus ligniperda* and *Hylastes ater* stridulations. Data points are averages from all 5 individuals. Beyond 40 cm species-specific stridulations become spectrally indistinguishable.

At distances up to 20 cm, all algorithms were able to accurately and automatically (> 97 %) discriminate stridulations of *H. ligniperda* from *H. ater* (Table II). After 40 cm, automatic identification reached chance levels, since the stridulations were so attenuated that they could not be discerned (Table II). This phenomenon can be visualized by plotting an ordination of amplitude, frequency, and time features in 2D space, where compact and segregated clusters are observable up to 20 cm for both species (Fig. 8). After 20 cm, the clusters became sparser until gradually merging at 60 cm, where stridulations of both species are embedded in the same subspace and cannot be discerned (Fig. 8).

**TABLE 2.**
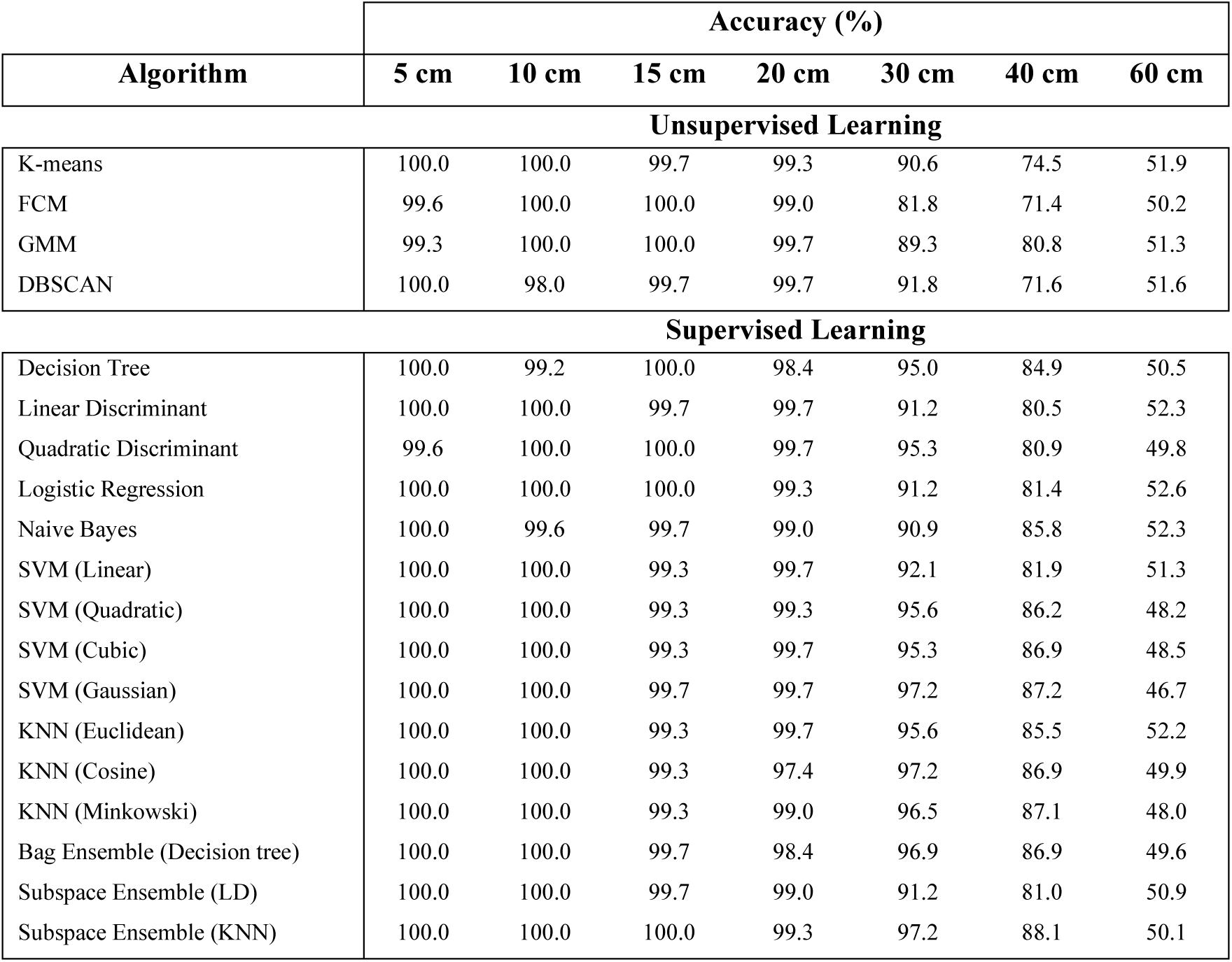
Accuracy results for several supervised and unsupervised machine learning approaches tested for a bi-class clustering/classification task: discriminating *Hylurgus ligniperda* and *Hylastes ater* stridulations at different distances.

| Algorithm | Accuracy (%) |  |  |  |  |  |  |
| --- | --- | --- | --- | --- | --- | --- | --- |
|  | 5 cm | 10 cm | 15 cm | 20 cm | 30 cm | 40 cm | 60 cm |
| **Unsupervised Learning** |  |  |  |  |  |  |  |
| K-means | 100.0 | 100.0 | 99.7 | 99.3 | 90.6 | 74.5 | 51.9 |
| FCM | 99.6 | 100.0 | 100.0 | 99.0 | 81.8 | 71.4 | 50.2 |
| GMM | 99.3 | 100.0 | 100.0 | 99.7 | 89.3 | 80.8 | 51.3 |
| DBSCAN | 100.0 | 98.0 | 99.7 | 99.7 | 91.8 | 71.6 | 51.6 |
| **Supervised Learning** |  |  |  |  |  |  |  |
| Decision Tree | 100.0 | 99.2 | 100.0 | 98.4 | 95.0 | 84.9 | 50.5 |
| Linear Discriminant | 100.0 | 100.0 | 99.7 | 99.7 | 91.2 | 80.5 | 52.3 |
| Quadratic Discriminant | 99.6 | 100.0 | 100.0 | 99.7 | 95.3 | 80.9 | 49.8 |
| Logistic Regression | 100.0 | 100.0 | 100.0 | 99.3 | 91.2 | 81.4 | 52.6 |
| Naive Bayes | 100.0 | 99.6 | 99.7 | 99.0 | 90.9 | 85.8 | 52.3 |
| SVM (Linear) | 100.0 | 100.0 | 99.3 | 99.7 | 92.1 | 81.9 | 51.3 |
| SVM (Quadratic) | 100.0 | 100.0 | 99.3 | 99.3 | 95.6 | 86.2 | 48.2 |
| SVM (Cubic) | 100.0 | 100.0 | 99.3 | 99.7 | 95.3 | 86.9 | 48.5 |
| SVM (Gaussian) | 100.0 | 100.0 | 99.7 | 99.7 | 97.2 | 87.2 | 46.7 |
| KNN (Euclidean) | 100.0 | 100.0 | 99.3 | 99.7 | 95.6 | 85.5 | 52.2 |
| KNN (Cosine) | 100.0 | 100.0 | 99.3 | 97.4 | 97.2 | 86.9 | 49.9 |
| KNN (Minkowski) | 100.0 | 100.0 | 99.3 | 99.0 | 96.5 | 87.1 | 48.0 |
| Bag Ensemble (Decision tree) | 100.0 | 100.0 | 99.7 | 98.4 | 96.9 | 86.9 | 49.6 |
| Subspace Ensemble (LD) | 100.0 | 100.0 | 99.7 | 99.0 | 91.2 | 81.0 | 50.9 |
| Subspace Ensemble (KNN) | 100.0 | 100.0 | 100.0 | 99.3 | 97.2 | 88.1 | 50.1 |

**FIG. 8.**
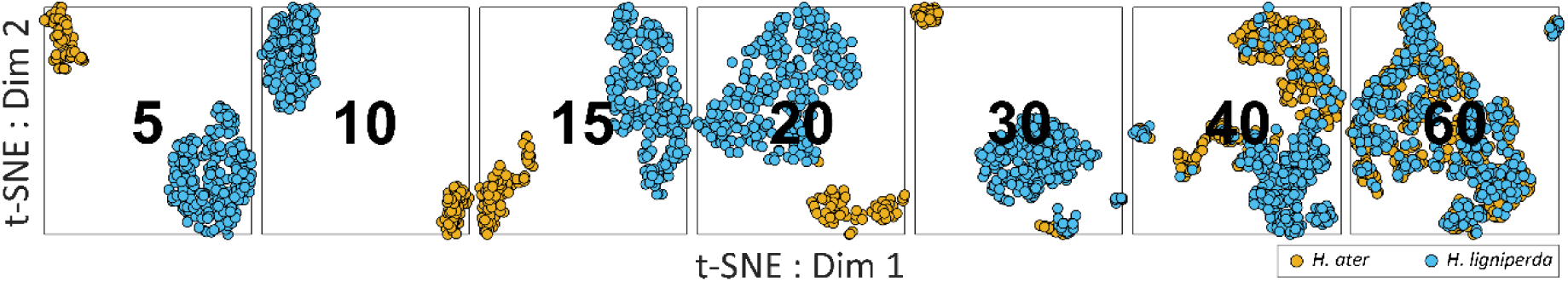
2D t-SNE visualization of individual stridulations of *Hylurgus ligniperda* (blue) and *Hylastes ater* (yellow) recorded at several distances (cm). Five acoustic features (mean amplitude, dominant frequency, centroid frequency, bandwidth, duration) were used for the ordination. Stridulations of both species become acoustically undistinguishable at 60 cm.

## 4. DISCUSSION

We characterized the propagation of stridulatory sounds of two bark beetle species (*H. ater* and *H. ligniperda*) through *P. radiata* logs, showing the effects of phloem and bark on signal attenuation over distance. We were able to correctly identify stridulatory sounds from insects of less than six mm length at distances of up to 40 cm. However, spectral content and signal amplitude attenuated with distance, particularly in the phloem tissue. Beyond 20 cm from the beetle, distance effects reduce the beetle signal bandwidth, which removes the part of the spectrum that allows species identification and makes *H. ligniperda* and *H. ater* stridulations difficult to distinguish. Nevertheless, the remaining content is sufficient to determine the presence of bark beetle activity after 40 cm, and additional temporal features may be used to tell species apart (e.g., call rate and inter-call interval), as these can be reliable species-specific descriptors (Bedoya *et al*., 2021).

In both species, power was concentrated between 3-7 kHz, which appears to be a general characteristic of Scolytinae (see supplementary material of Bedoya *et al*., 2021). We found that 4-6 kHz, where most of the energy is concentrated, was the optimal frequency band to detect stridulations. This frequency is also one of the least attenuated by pine trees (Price *et al*., 1988). Our results concord with previous experimental models for the propagation of sound inside wood (Burns, 1979; Price *et al*., 1988), suggesting an acoustic impedance matching between the beetle stridulatory mechanism and the medium. Measurements of sound speed in *Pinus radiata* have been previously reported for logs and standing trees from New Zealand forests (Wang *et al*., 2006); consequently, these measurements were not part of our experimental design. The mean sound speed in *P. radiate* is 2277±496.1 m/s (mean±SD) for standing trees and 2120±363.5 m/s for logs of 3.66 m long (Wang *et al*., 2006). Accurate descriptions of the dependence of sound speed in *P. radiata* on tree age, type of tissue, length and width of the tree, and moisture content are described in Grabianowski *et al*. (2006), Wang *et al*. (2006), and Toulmin *et al*. (2007).

We also propose a series of exponential models for the power decay of the stridulations depending on beetle species, type of tissue, and distance. Removing the bark did not significantly reduce the signal power, suggesting that beetles can be accurately detected without removing the bark. Furthermore, our machine learning analyses suggest that species can be reliably identified (> 97% accuracy) at short distances of < 20 cm, and with relatively good accuracy (> 70%) up to 40 cm. After 40 cm, our experimental model shows that most of the energy has already dissipated, and none of the tested clustering or classification algorithms was able to provide accurate identification results. Nonetheless, the presence of bark beetles can still be detected at further distances in the 4-6 kHz frequency band if the species is large enough (e.g., *H. ligniperda*). The accuracies obtained using supervised and unsupervised approaches are almost identical up to 20 cm, from 30 cm onwards supervised learning techniques become advantageous. However, the increase in accuracy is not large enough to overcome the benefits of unsupervised learning (e.g., no need for data labelling).

As bark beetles are amongst the smallest of the woodborers (Kirkendall *et al*., 2015), for bigger taxa, such as some pinhole borers (Platypodinae), which tend to generate louder stridulations than most bark beetles (Bedoya *et al*., 2021), we would expect similar attenuation patterns and longer detectability ranges than those found here. The smallest woodborer with acoustic communication capabilities (*Ips avulsus*, 2.5 mm) is relatively loud for its size and has a similar amplitude range to *H. ater* (Bedoya *et al*., 2021). Consequently, we estimate that deploying an array of sensors spaced at distances of 40 cm should be enough to detect stridulations of any bark-or wood boring species in logs similar to those of *P. radiata*. The key remaining issue for the detection of the potential presence of such insects is how to elicit *ad libitum* sound production under the bark of trees, so that the stridulations can be detected in a species-specific manner to identify the presence of woodborers in logs and standing trees. Chemical, acoustic, and luminous stimuli can elicit acoustic communication in several species (Rudinsky and Michael 1972; Hofstetter *et al*., 2019; Bedoya *et al*., 2019b). However, integrating these stimuli with acoustic detection and identification methods has yet to be addressed, especially when the target organism is hidden under the bark.

For acoustic identification purposes, deploying an array of sensors 20 cm apart is enough to detect and identify a species. At distances below 20 cm, between the source and the sensor, the spectral content of the stridulation does not change enough to make the species indistinguishable. Increasing the distance between sensors may increase the detectability range, but may affect the accuracy of the species identification. Depending on the application and the need for accuracy, 40 cm is a good compromise, as most stridulations are still detectable and the identification accuracy is above 70%. If the acoustic identification set-up is located in an environment with much background noise and the frequency range needs to be restricted, 4 to 6 kHz is a useful band to analyze, as this is where most of the energy is concentrated. Bark beetles live in the phloem, but part of their bodies are usually in contact with the bark tissue, generating a direct coupling with the drier outermost bark layer. Consequently, from the attenuation standpoint, piercing the tree in order to place the sensor in the phloem layer does not appear to provide a substantial benefit, as stridulations attain similar detectability ranges in both types of tissues. Bark is the most accessible contact point between the sensor and the tree stem; thus, placing the sensor on the bark surface of a tree or stem does not jeopardize species detection and does not produce tissue damage.

No studies have been performed on tree soundscapes or acoustic interactions of bark-or wood boring beetles in their natural habitat, despite the prevalence of acoustic activity in insects living inside trees. Our study provides a better understanding of the propagation of stridulatory under the bark of trees and contributes towards the development of techniques to study bark- and woodborers in nature. We provide information on how these beetles could be acoustically detected and identified, where to position sensors, and in which part of the frequency of the acoustic spectrum to search for identifying information. We hope this study promotes understanding of acoustic communication inside tree tissues and its role in animal interactions. An appreciation of how stridulatory signals propagate inside tree tissues should aid in our understanding of colonization patterns, gallery structure, and niche-partitioning between cohabitating species. We also hope this work establishes new ground for technological development to aid in automatic acoustic detection approaches for biosecurity purposes. As some bark beetles are of significant economic and biosecurity importance (McCarthy *et al*., 2013; Grégoire *et al*., 2015), a clear understanding of acoustic signal propagation through bark and wood can enhance efforts to determine the presence and species identity of potential pest species at borders.

## DATA ACCESSIBILITY

Acoustic datasets and algorithms used during this research are available at:

Data: https://doi.org/10.6084/m9.figshare.19233087.v1

Code: https://github.com/carolbedoya/Beetle-Sounds-Inside-Wood

## ACKNOWLEDGEMENTS

This project was supported by the New Zealand Ministry of Business, Innovation, and Employment (MBIE), grant C04X1407, the Better Border Biosecurity Collaboration (b3nz.org) via MBIE Core Funding to Scion, and Catalyst: Seeding funding from the Royal Society of New Zealand (grant CSG-FRI1701).

